# An operational methodology to identify Critical Ecosystem Areas to help nations achieve the Kunming-Montreal Global Biodiversity Framework

**DOI:** 10.1101/2023.05.03.539215

**Authors:** Ruben Venegas-Li, Hedley S. Grantham, Hugo Rainey, Alex Diment, Robert Tizard, James E.M. Watson

**Affiliations:** School of Earth and Environmental Sciences, The University of Queensland, Brisbane, Qld, Australia; Centre for Ecosystem Science, University of New South Wales, Sydney, NSW, Australia; Bush Heritage Australia, Melbourne, Victoria, Australia; Wildlife Conservation Society, 2300 Southern Boulevard, Bronx 10460, NY, USA

## Abstract

The Kunming-Montreal Global Biodiversity Framework (GBF) will become the most important multilateral agreement to guide biodiversity conservation actions globally over the coming decades. An ecosystem goal and various targets for maintaining integrity, restoring degraded ecosystems, and achieving representation in conservation areas feature throughout the GBF. Here, we propose an operational framework that combines disparate information on ecosystem type, extent, integrity, levels of protection, and risk of collapse to support the identification of irreplaceable ‘Critical Ecosystem Areas’ (CEAs), to help advance these ecosystem targets. The framework classifies each component ecosystem based on its integrity, importance in ensuring no ecosystem collapse and its relative value to achieving representation if protected. These CEAs are immediate conservation opportunities, given that they achieve multiple ecosystem goals and targets in the GBF. We showcase its application using Myanmar’s forested ecosystems as a case study and argue that it could be immediately used across all terrestrial ecosystems.

## Introduction

The Kunming-Montreal Global Biodiversity Framework (GBF) (CBD 2022) has targeted sustaining and enhancing ecosystem area, connectivity, resilience, and integrity at the forefront of its vision, goals (Goal A), and Targets (1, 2, 3, 12). If implemented by Convention for Biological Diversity (CBD) signatory nations, they will likely be a core plank in efforts to advance all biodiversity conservation agendas successfully, considering that functioning, resilient ecosystems are essential for sustaining species and genetic diversity (Di Marco et al. 2018; Watson et al. 2020; Nicholson et al. 2021). It is now well established that ecosystem degradation increases species extinction risk (Barlow et al. 2016; Betts et al. 2017, 2022), reduces the capacity to sustain essential ecosystem functions and services (IPBES 2019), and diminishes overall resilience to climate change (Watson et al. 2018; Pörtner et al. 2021). Thus, meeting this ecosystem goal is also central to meeting the GBF’s Goals B and C for enhancing nature’s contributions to people and sharing the benefits of genetic diversity fairly and equitably. Moreover, focusing on ecosystem conservation and restoration will help advance other global agendas, such as abating the impacts of climate change on biodiversity and vulnerable human communities (Martin & Watson 2016; IPCC 2023).

The key components for effectively implementing the GBF’s Goal A must include assessing and planning for ecosystem type, extent, integrity, and risk of collapse (Nicholson et al. 2021), and ensuring that representative samples of all ecosystems exist within conservation areas (Jetz et al. 2021). There are now widely accepted practical definitions of ecosystem types, risk of collapse, and integrity (or related concepts of condition and degradation), all formalized in global standards via efforts like the IUCN Red List of Ecosystems (RLE) (Keith et al. 2015) and the UN System for Environmental Economic-Accounting (King et al. 2021). Moreover, data on all these ecosystem components are largely available via the many global, national, and regional ecosystem maps and RLE assessments (Nicholson et al. 2021, iucnrle.org); and the availability of these data is expected to improve quickly (Hansen et al. 2021). However, a significant shortfall exists in successfully incorporating these ecosystem components coherently to support spatial planning for ecosystem conservation, and necessary for achieving GBF’s Targets 1 and 14.

The Kunming-Montreal GBF provides a mandate to signatory countries of the CBD and civil society for implementing strategies that will allow the global community to meet its vision of ‘living in harmony with nature’. Prompt action is needed to ensure effective implementation, given the present rates of biodiversity loss and ecosystem degradation (Diaz et al. 2019) and the proximity of 2030, a year in which several action-oriented targets should be met. Thus, our aim here is to propose an operational framework that integrates information on ecosystem type, extent, degree of integrity and risk of collapse, which can then inform spatial action planning efforts to prioritise ecosystem-based efforts needed to advance the Kunming-Montreal GBF agenda. Specifically, this framework identifies key ecosystem areas, which we name Critical Ecosystem Areas (CEAs), which are of utmost importance for immediate conservation attention as their protection is needed to advance towards achieving the goals and targets on ecosystem representation, integrity, and risk of collapse.

Using Myanmar forested ecosystems as a case study, we illustrate how CEAs can be identified and can inform spatial planning towards achieving various of the GBF’s 2030 action targets. Specifically, their identification is essential for the retention of intact areas (Target 1), ecosystem restoration (Target 2), and protected areas under the ‘30 × 30 vision’ (Target 3). CEAs also contribute to the sustainable use of ecosystems (Goal B), for example, by informing the design and application of measures across the mitigation hierarchy (Jones et al. 2022), which can support achieving Targets 14 and 15. Notably, the existing methods and data we use can be easily adapted in the framework, making it implementable by any nation - and other stakeholders-immediately.

## Methods

### Operational Framework

Our framework (Fig 1) integrates information on ecosystem type, extent, integrity, risk of collapse and degree of protection using a spatial planning approach to identify CEAs. CEAs should be considered a priority for conservation actions based on their importance to achieving the GBF’s ecosystem-related goal and targets. First, the framework subdivides ecosystems into subunits to assess their relative importance for achieving ecosystem representation targets in areas with the highest possible integrity, while considering broad landscape scale connectivity and in relation to existing protected areas. Here, particular attention can be placed on ecosystems at higher risk of collapse by setting higher representation targets for these (recognising that representation targets could be country-specific (Maron et al. 2019)). Next, the map of relative importance for conservation is combined with the ecosystem integrity map to produce a classification of ecosystems that can inform conservation actions based on an area’s potential contribution to achieving the set goals, and its integrity. We provide a case study applying this methodology using Myanmar forest ecosystems.

**Figure 1.**
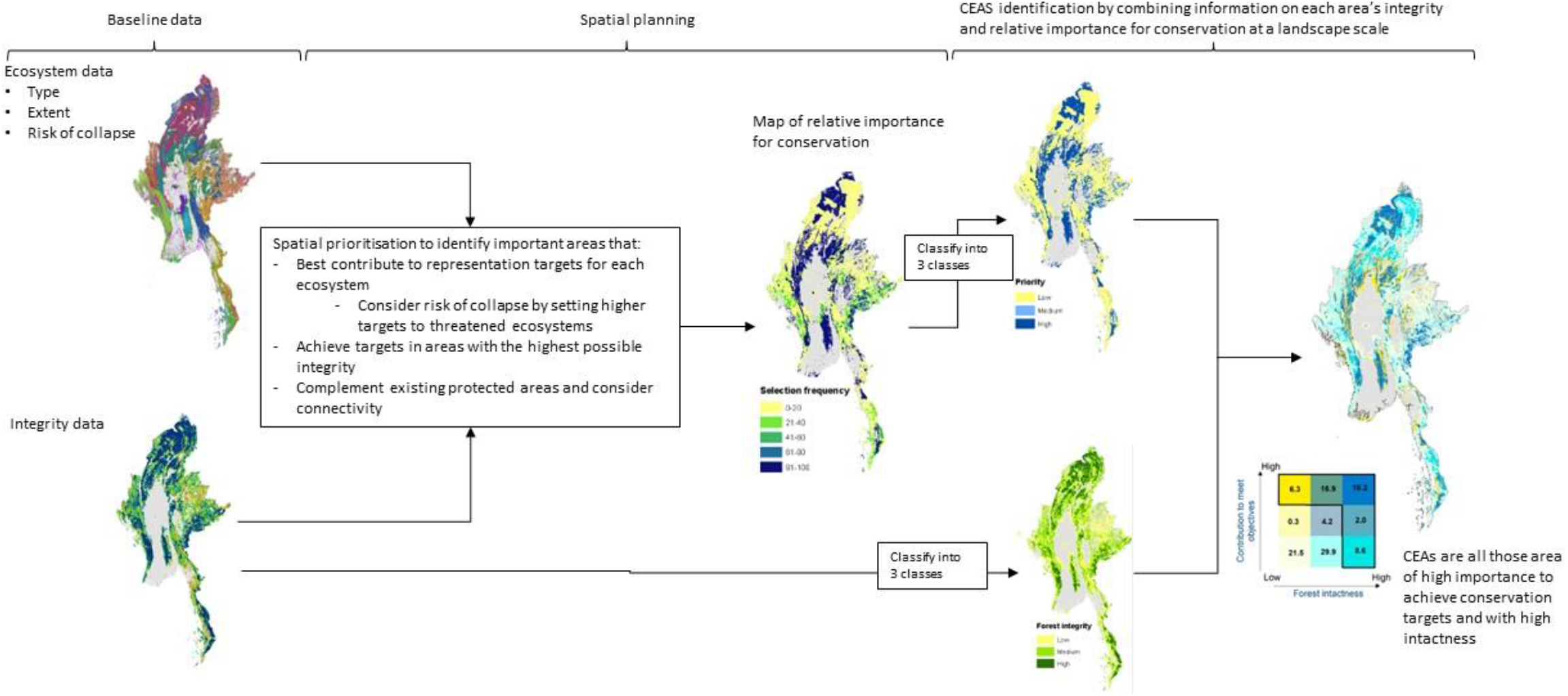
Flowchart of a methodological approach that combines ecosystem integrity, extent, and risk of collapse using a spatial prioritisation analysis to produce a spatial tool that can inform ecosystem conservation actions towards achieving the Global Biodiversity Framework’s ecosystem milestones and goals.

### Myanmar’s case study

#### Ecosystem data and level of protection

We used a dataset of natural ecosystems of Myanmar (Murray et al. 2020b, 2020a), with information on ecosystem type, extent, and risk of collapse mapped at 90 m spatial resolution. We restricted our analysis to 46 mapped forest ecosystems, as we only have integrity data for forest ecosystems. We used the forest landscape integrity index (FLII) (Grantham et al. 2020), a continuous index of forest condition determined by the expected degree of human modification. The index integrates data on observed human pressures (e.g., roads and canopy loss), inferred pressures associated with people (e.g., infrastructure effects), diffuse processes (e.g., increased access to hunting and logging), and anthropogenic changes in forest connectivity. The FLII was mapped globally at a 300m spatial resolution, using the forest cover for the start of 2019 as a baseline for forest ecosystems, and scaled between 0 (low integrity) and 10 (high integrity). We determined the level of protection for each forested ecosystem using protected area data curated by Murray et al. (2020a).

### Spatial prioritisation: identifying relative importance to achieve conservation targets

We used the decision support software Marxan (Possingham et al. 2000) to assess the relative importance of a given forest area to achieve ecosystem representation targets in places in the best possible condition. Marxan uses a simulated annealing algorithm to identify multiple near-optimal configurations of sites in a study region where defined conservation targets can be achieved while minimising cost (Ball et al. 2009). Here, we subdivided the forest landscape into 4 km^2^ units, henceforth called planning units.

We set as a target in our analysis to represent at least 30% of the extent of each ecosystem in a suite of planning units, aligning the targets with the 30 × 30 vision. However, to highlight the importance of conserving ecosystems at high risk of collapse, we set a target of 100% for the 13 critically endangered and endangered ecosystem types. We also predetermined into all the solutions the existing protected area. Thus, threatened ecosystems and protected areas are pre-emptively considered as highly important areas for conservation (and thus CEAs), and newly selected areas will be complementary and connected to these. To achieve a degree of connectivity between selected sites in Marxan, we calibrated the boundary length modifier following Stewart and Possingham (2005).

To achieve representation targets in areas in the best possible condition, we used the inverse value of the mean FLII for each planning unit as a cost in Marxan. Thus, high-integrity areas represent a low cost, and will be selected where possible to minimise the solution’s total cost.

Marxan provides only near-optimal solutions; thus, different planning unit arrangements can result when Marxan is run multiple times. We ran Marxan 100 times with 50,000,000 iterations per run to obtain a selection frequency for each planning unit. The selection frequency shows the number of times a particular planning unit was selected across all 100 runs and provides a measure of their relative importance in achieving the set objectives.

### Critical Ecosystem Areas and implementation examples

To obtain a map showing for each planning unit both its relative contribution to ecosystem conservation objectives and its condition, we combined the selection frequency with the ecosystem integrity map. We first reclassified each map into three classes (low, medium, and high). The selection frequency map classes were classified as low (0-40), medium (≥ 40 and <70), and high (≥ 70). The integrity map was classified into three classes following Grantham et al.(Grantham et al. 2020) : low (values < 6), medium (≥ 6 and <9.6), and high forest integrity (≥ 9.6). Combining the maps results in a bivariate map with nine classes. We have called those areas with high relative conservation value and/or high integrity as Critical Ecosystem Areas (CEAs), arguably the areas that will better contribute towards achieving the GBF’s ecosystem goals and targets.

We illustrate with three examples how this framework can inform conservation and sustainable use planning that contributes to implementing the GBF. First, we overlayed the resulting CEAs with the protected areas data to quantify how much of their area is unprotected and show how these data can inform protected areas planning. We then quantified the area of CEAs based on their integrity, to illustrate how this methodology could inform decision-making around application of the mitigation hierarchy (Phalan et al. 2018; Jones et al. 2022) for development planning, specifically avoidance of development in the most important sites with highest integrity, and how restoration efforts, including compensatory offsets, could be located in areas with lower integrity. Finally, we overlayed our 9-class map with hypothetical agricultural concessions in a region of Myanmar, to illustrate how these can be used to report both the individual and the cumulative impact of single or multiple development projects in the landscape.

## Results

Of the 46 forest ecosystems in Myanmar, 13 were classified as either Critically Endangered or Endangered (Fig 2, Table S1). While the mean integrity of all 1 km^2^ forest pixels in the country is 7.1 ± 2.8, the range of values between and within ecosystems varies widely (Fig 2) with some forested ecosystems (e.g., Tanintharyi cloud forest) having few areas left that can be considered high integrity. Approximately 28% of the country’s remaining forest has low integrity, 51% has medium integrity, and 21% has high integrity.

**Figure 2.**
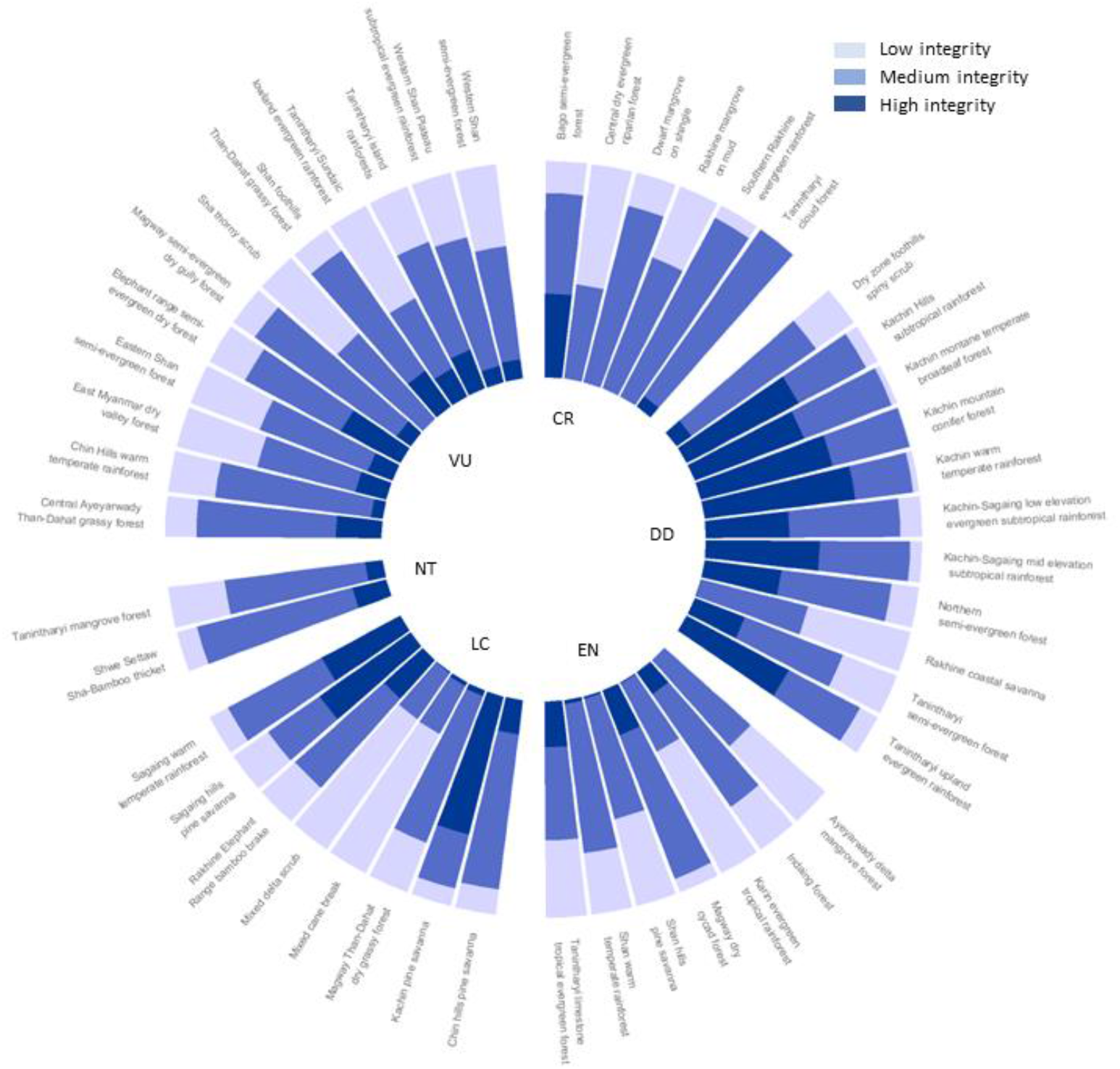
Myanmar’s forest ecosystem types and the proportion of their distribution that has low, medium, and high forest integrity according to the Forest Landscape Integrity Index (Grantham et al. 2020). Forest types are grouped by risk of collapse status as per a Red List of Ecosystem assessment (CR= Critically endangered, EN= Endangered, VU= Vulnerable, NT= Near Threatened, DD= Data Deficient, LC= Least Concern).

A 24% of Myanmar’s forested landscape is essential for conservation based on either its status as a protected area (9.5%) or because it corresponds with ecosystems at a very high risk of collapse (15.5%). Through the spatial prioritisation analysis, we identified an additional 9.4% of forested landscape as having a relatively high importance for conservation (selection frequency ≥ 70), representing areas with the highest possible integrity and complementary to the already predefined important areas (Figure 3a, 3b).

**Figure 3.**
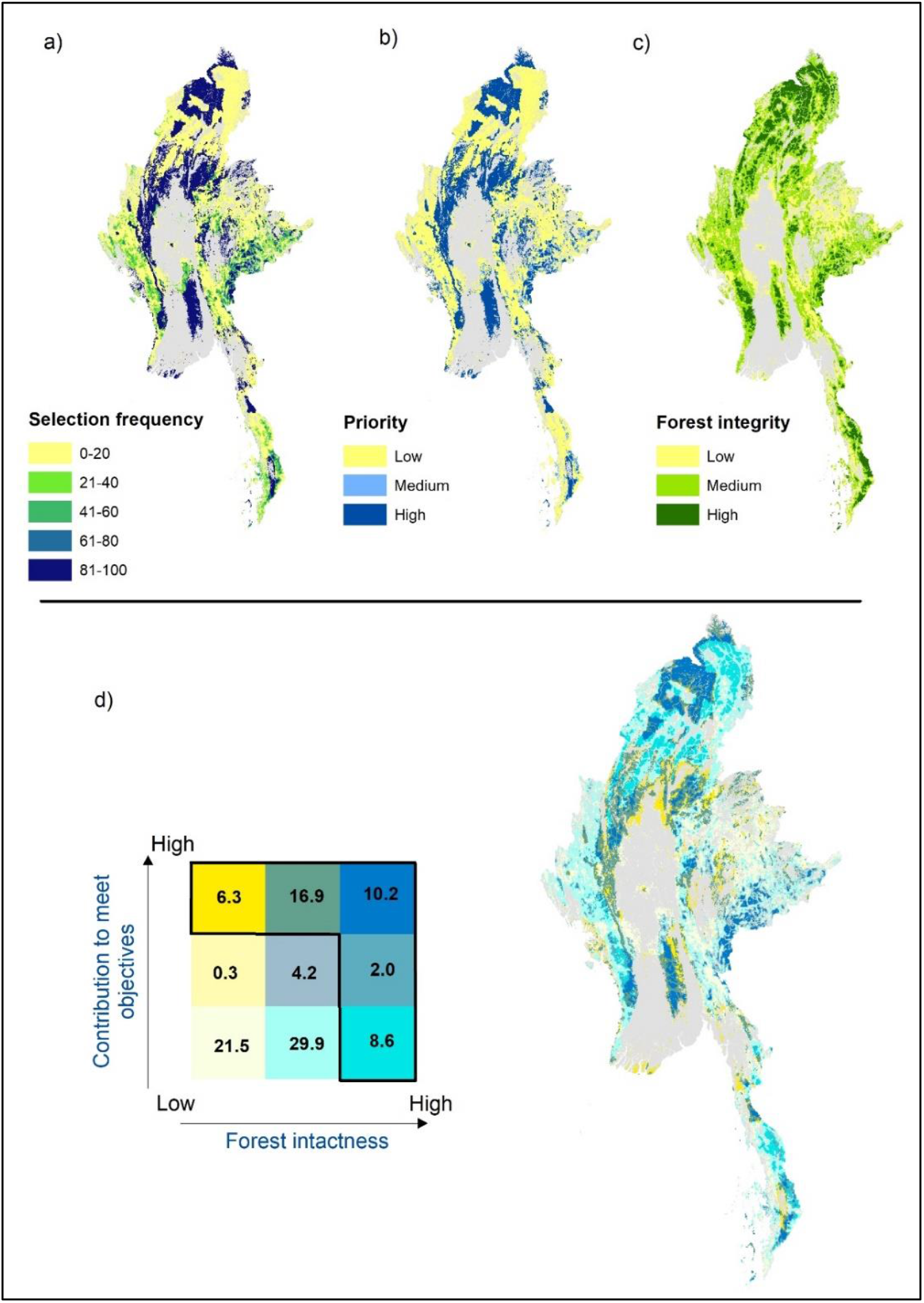
Classification of forest ecosystems of Myanmar based on their relative importance to achieve conservation objectives and their integrity. A spatial conservation prioritisation analysis that combines ecosystem type, extent, risk of collapse, and integrity results in map of relative importance for planning sites to achieve conservation objectives (a). This map of priorities and the map of forest integrity are reclassified into 3 classes, low, medium, and high (b and c respectively). Finally, the reclassified maps are combined to classify each forest grid cell into one of 9 classes based on their relative importance to contribute to conservation objectives, and on their integrity. The number inside the legend for (d) corresponds to the percentage area that each category occupies within the landscape; the five categories bounded by the black border lines correspond to what we have termed Critical Ecosystem Areas.

Combining the maps of relative importance to achieve conservation objectives and the map of ecosystem integrity (Fig 3b, 3c) produced a metric that classifies each forest pixel into one of nine different classes (Fig 3d). Each of these classes indicates how important a particular pixel is for achieving conservation objectives for ecosystems, as well as its integrity. Our results show that approximately one-fifth of Myanmar has both low forest integrity and is unlikely to contribute to meeting the conservation objectives. In contrast, at least 44% of the forest landscape (27% of the country) can be considered a Critical Ecosystem Area (Fig 4a), i.e., these are areas with high relative importance to achieving ecosystem conservation, high ecosystem integrity, or both.

**Figure 4.**
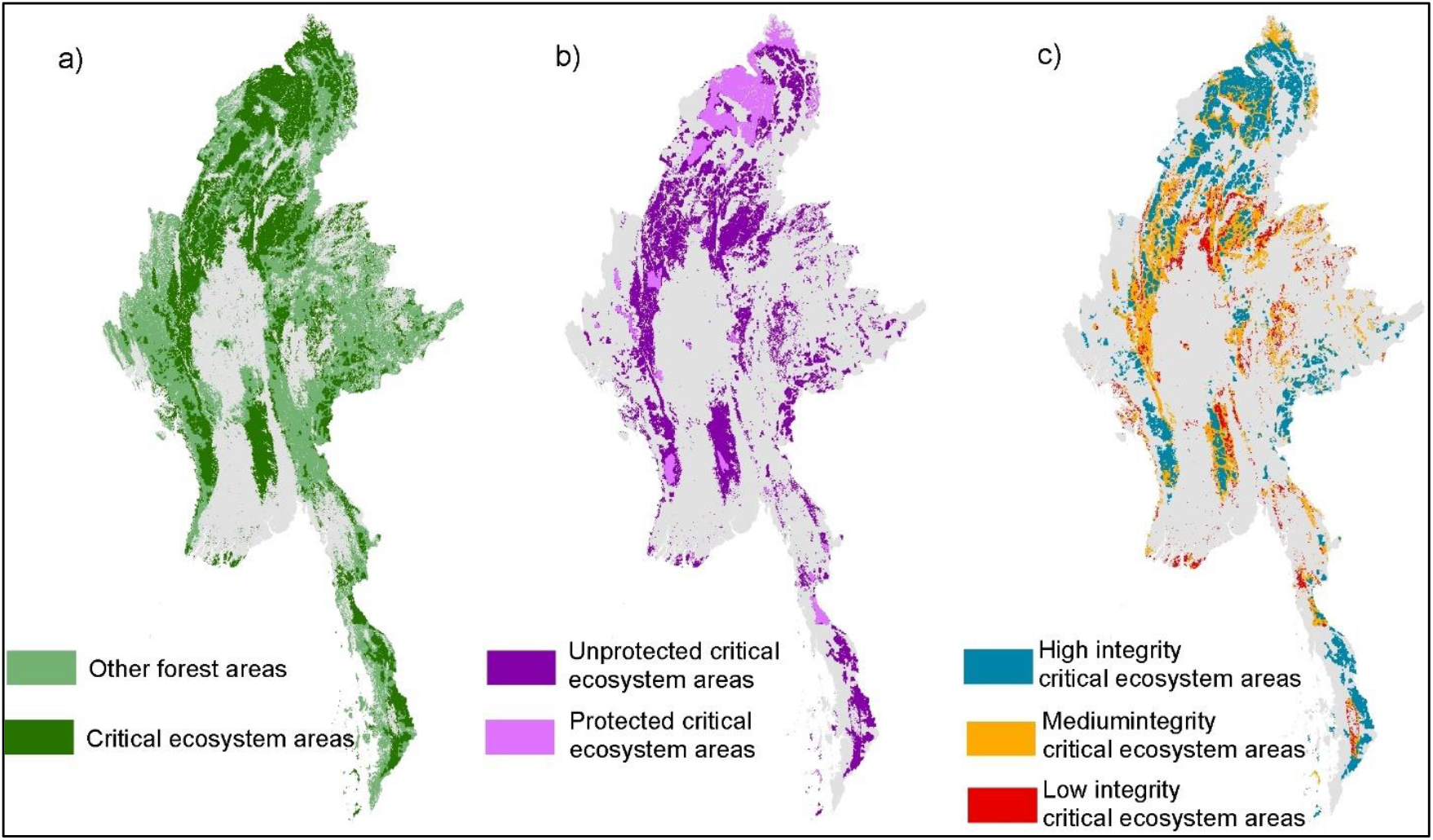
Identifying Critical Ecosystem Areas (a) which can be used to inform conservation actions planning such as protected area expansion (b) and restoration efforts in CEAs with medium and low ecosystem integrity (c).

We illustrated potential ways our framework could be used to inform different types of spatial conservation planning efforts (Fig 4). For example, only 20.4% of the areas we identified here as CEAs are part of the current Protected Area System of Myanmar (4b and Table 1). Future expansion of this PA system should consider the remaining 79.6% of unprotected CEAs as prime candidate areas for an expansion aiming to contribute the most towards the GBF’s ecosystem goal and targets. Another application where our framework could result useful is in planning of restoration efforts (Fig 4b). Here we show that 52.8%of CEAs are somewhat degraded (low and medium integrity) and could be considered as a high priority for forest restoration efforts.

**Table 1.** Critical Ecosystem Areas and their overlap with protected areas.

| Integrity | Area (km<sup>2</sup>) | % of CEAs area | % of Myanmar's terrestrial area | Area protected (km<sup>2</sup>) | CEAs area protected (%) | CEAs area not protected (%) |
| --- | --- | --- | --- | --- | --- | --- |
| High | 86,531 | 47.3 | 12.9 | 21,215 | 24.5 | 75.5 |
| Medium | 70,222 | 38.4 | 10.5 | 14,818 | 21.1 | 78.9 |
| Low | 26,294 | 14.4 | 3.9 | 1,356 | 5.2 | 94.8 |
| Total | 183,047 | 100 | 27 | 37,389 | 20.4 | 79.6 |

Our framework and the resulting classification system could be used in development planning to inform avoidance of impacts on CEAs. For example, 21.5% of the forest has low integrity and importance towards achieving conservation objectives (Fig 3d) and could be considered as suitable areas where development leading to forest degradation could occur, if offset safeguards are in place. However, CEAs should be avoided as they are irreplaceable and not offsetable. This showcases the potential use of the data generated by our framework as a reporting tool (Table 2) to understand the risk that a particular development project or group of projects could pose to biodiversity and guide the application of the mitigation hierarchy based on a place’s integrity and importance to achieve ecosystem objectives. In our example, the overlap of agricultural concessions in southern Myanmar’s forests (and identified CEAs) is considerable (Table 2), clearly highlighting the risk of different ecosystem types to clearing from a single sector.

**Table 2.** Overlap between Myanmar’s forest ecosystems with agricultural concessions, categorised based on their relative importance for conservation and integrity. The resulting classification from the CEAs framework can be used as a reporting tool for businesses and countries in terms of understanding the risk that single or multiple activities pose to ecosystems. Numbers in bold represent the area (km^2^) of each category that agricultural concessions would impact if they went forward.

| Ecosystem | Critical ecosystem areas (km<sup>2</sup>) |  |  | Other areas (km<sup>2</sup>) |  |
| --- | --- | --- | --- | --- | --- |
|  | High FLII | Medium FLII | Low FLII | Medium FLII | Low FLII |
| East Myanmar dry valley forest | 3,746 | 8,741 | 2,376<br>**8** | 5,545<br>**1** | 13,709<br>**1** |
| Mixed delta scrub | 1 | 39 | 227<br>**3** | 7<br>**2** | 538<br>**9** |
| Tanintharyi limestone tropical evergreen forest | 366<br>**1** | 819<br>**60** | 767<br>**96** | 0<br>**0** | 4<br>**0** |
| Tanintharyi mangrove forest | 197 | 1169 | 361 | 779<br>**74** | 653<br>**9** |
| Tanintharyi semi-evergreen forest | 4727<br>**18** | 5786<br>**2** | 1669<br>**4** | 4123<br>**39** | 4310<br>**38** |
| Tanintharyi Sundaic lowland evergreen rainforest | 905<br>**28** | 1598<br>**1** | 1477<br>**7** | 898<br>**371** | 2336<br>**200** |
| Tanintharyi upland evergreen rainforest | 3962<br>**42** | 2364<br>**5** | 482<br>**7** | 588<br>**133** | 384<br>**28** |

## Discussion

The Kunming-Montreal GBF is now the most important multilateral agreement to guide global biodiversity conservation actions for the next three decades. The final text of the GBF includes a core ecosystem component as part of its Goal A and several ecosystem-based targets (Targets 1,2, and 12), which are currently treated independently (Nicholson et al. 2021). Here, we integrated the key components of the ecosystem goal and targets into an operational framework to support the GBF implementation. The framework has three key characteristics that make it highly relevant for conservation and development planning and the GBF implementation in particular: i) it allows identifying irreplaceable ecosystem areas and measurement of ecosystem integrity, here called Critical Ecosystem Areas (Goal A, Targets 1, 2); ii) it can be implemented immediately across ecosystems and countries; and iii) it simplifies the complexity of the landscape integrity matrix to facilitate spatial planning (Targets 1, 2, 3) and application of the mitigation hierarchy (Targets 14 and 15), particularly for complex cumulative impacts.

Critical Ecosystem Areas represent samples of each ecosystem with the highest integrity, complementary to existing protected areas and that maintain some degree of connectivity throughout the landscape. CEAs also capture those ecosystems most threatened with collapse and remaining intact areas. As all these are critical components of the GBF, CEAs should be considered irreplaceable areas that must be retained to achieve the GBF’s goals and targets. For example, Target 1 calls to bring the loss of areas of high biodiversity importance such as ecosystems of high integrity close to zero by 2030, only seven years from now. The loss of any CEAs would make it almost impossible to achieve this target. As representatives of the highest integrity within their ecosystems CEAs hold high values such as biodiversity and carbon in the case of forests (Pörtner et al. 2021); recovering any of these values if lost or degraded will take decades to centuries (Jones et al. 2018; Watson et al. 2018), incompatible with the GBF timeframes. We argue that any industrial development in CEAs should be avoided as much as possible; losses of these areas are unlikely to be offsetable, because identifying equivalent ecological benefit elsewhere is likely impossible, and their restoration would be technically and politically complex and will not occur in short time frames (Gibbons et al. 2016; Sonter et al. 2020), or might not occur at all (Lindenmayer et al. 2017).

The framework is implementable in the near term, given that the data is available for a considerable number of nations and ecosystems, which responds to the GBF’s call in the 2030 Milestones to take urgent action, and highly relevant to conservation practice in the context of the present biodiversity crisis (Leclère et al. 2020). For example, approximately 60 countries have completed their national Red List of Ecosystem assessments, and 20 countries have subsets of ecosystems such as forests assessed. A global RLE for terrestrial ecosystems may be finished by 2025 (iucnrle.org) at a minimum, the CEAs framework could be immediately implemented for most of the World’s forest ecosystems, which cover approximately one-third of Earth’s terrestrial area (FAO & UNEP 2020). The next challenge will be to implement this framework in other freshwater, marine, and non-forest terrestrial ecosystems.

### Simplifying landscape matrix complexity

Classifying CEAs based on their relative conservation importance, ability to achieve representation targets, and overall integrity allows for an important simplification of the complexity around disparate ecosystem goals and agendas. The framework responds to Target 1 of the GBF, ensuring that all land and sea areas globally are under spatial planning. CEAs identified through the framework can directly inform implementation efforts around achieving other action targets as we illustrated here, for example ensuring the restoration of at least 30% of degraded ecosystems (Target 2) and ensuring that 30% of land and sea areas are under protected area and other effective area-based conservation measures (Target 3). CEAs conservation will ensure the achievement of the ecosystem-based components of Targets 1-3, it will also indirectly contribute to advancing other targets related to reducing species extinction (Target 4), reducing threats to biodiversity (e.g., Target 8), and meeting peoples’ needs through sustainable use and benefit sharing (Targets 11, 14-15). CEAs can also be used to inform the planning, design, and application of measures across the mitigation hierarchy by businesses, countries, and financial lenders (Targets 14-15). For example, identifying CEAs can be used in government spatial planning for development and the early phases of project design to inform particularly the avoidance stage (Jones et al. 2022), or for supporting business and countries to report their impacts on biodiversity (Target 15). As CEAs are created at a landscape scale, they can be used to consider the cumulative impacts of multiple development projects on ecosystems (Franks et al. 2010; Whitehead et al. 2017) and avoid them through better planning (Target 14), as well as providing a more accurate accounting towards losses and gains of biodiversity.

## Conclusion

Here we integrate the core concepts of an ecosystem-based conservation goal under an operational framework to support the implementation of the Kunming-Montreal GBF, applicable to all forested countries in the near-term. We argue this framework directly responds and could support operationalizing targets for achieving spatial planning and retention of intact areas (Target 1), ecosystem restoration (Target 2), protected area planning (Target 3), and supporting better decision making around the mitigation hierarchy (Targets 14 and 15) including actions to avoid impact on the most important sites. By mapping CEAs, it will enable nations to better measurement of progress towards integrity goal and targets which will support better decision-making around the mitigation hierarchy, including actions to avoid impacts from development on the most important sites, restoration of sites with low integrity and identification of new protected areas. The framework is flexible, and it can be adapted to the varying spatial conservation priorities that can be considered in a national application.

## Acknowledgements and Data

Alcoa Foundation funded development of this methodology and part of RVL’s time. RVL’s time was also funded by University of Queensland through a strategic grant. Data and results can be accessed through The University of Queensland’s e-space.

